# Identification and Structure-Activity Relationship of HDAC6 Zinc-finger Ubiquitin Binding Domain Inhibitors

**DOI:** 10.1101/268557

**Authors:** Renato Ferreira de Freitas, Rachel J. Harding, Ivan Franzoni, Mani Ravichandran, Mandeep K. Mann, Hui Ouyang, Mark Lautens, Vjayaratnam Santhakumar, Cheryl H. Arrowsmith, Matthieu Schapira

## Abstract

HDAC6 plays a central role in the recruitment of protein aggregates for lysosomal degradation, and is a promising target for combination therapy with proteasome inhibitors in multiple myeloma. Pharmacologically displacing ubiquitin from the zinc-finger ubiquitin-binding domain (ZnF-UBD) of HDAC6 is an underexplored alternative to catalytic inhibition. Here, we present the discovery of a HDAC6 ZnF-UBD-focused chemical series and its progression from virtual screening hits to low micromolar inhibitors. A carboxylate mimicking the C-terminal extremity of ubiquitin, and an extended aromatic system stacking with W1182 and R1155 are necessary for activity. One of the compounds induced a conformational remodeling of the binding site where the primary binding pocket opens-up onto a ligand-able secondary pocket that may be exploited to increase potency. The preliminary structure-activity relationship accompanied by nine crystal structures should enable further optimization into a chemical probe to investigate the merit of targeting the ZnF-UBD of HDAC6 in multiple myeloma and other diseases.

## Introduction

HDAC6 is the main cytoplasmic deacetylase in mammalian cells, and the only deacetylase containing a zinc finger ubiquitin-binding domain (ZnF-UBD).^1–3^ Targets deacetylated by HDAC6 include α-tubulin^4^, HSP90^5^, and cortactin^6^ among others.^7,8^ HDAC6 is involved in a diverse array of biological processes, including cell motility^9^, immune synapse formation^10^, viral infection^11^, and the degradation of misfolded proteins^12^, and has been associated with diseases.^13^ In particular, HDAC6 recruits polyubiquitinated protein aggregates via its ZnF-UBD, and loads misfolded proteins onto dynein for microtubule-guided transport to the lysosome.^12^ This lysosomal degradation pathway is an alternative mechanism to proteasomal degradation for the clearance of misfolded proteins^12^, and chemical inhibition of HDAC6 represents a promising strategy to overcome resistance to proteasome inhibitors in multiple myeloma and lymphoma.^14–16^

Current HDAC6 inhibitors are targeting the catalytic domain of the enzyme, thereby inhibiting the de-acetylation of microtubules, and antagonizing the transport of protein aggregates to the aggresome^17^. An alternative mechanism of action is to target the ZnF-UBD domain of HDAC6, which would antagonize HDAC6-mediated recruitment of ubiquitinated protein aggregates to microtubules. We have recently developed a collection of assays to enable the development of HDAC6 ZnF-UBD chemical probes, and reported fragment hits occupying its binding pocket.^18^ Here, we describe the design and discovery of a low micromolar HDAC6 ZnF-UBD inhibitor guided by virtual screening and X-ray crystallography. The emerging structure-activity relationship (SAR) defines several scaffolds as attractive starting points for further elaboration into high affinity inhibitors.

### Design

The crystal structure of the HDAC6 ZnF-UBD was previously solved in the apo form (PDB: 3C5K), and in complex with an ubiquitin C-terminal pentapeptide, RLRGG, (PDB: 3GV4) as well as in complex with full-length ubiquitin (PDB: 3PHD).^19^ The C-terminal carboxylate of the ubiquitin peptide binds in a deep cavity of HDAC6. Binding is stabilized by an extensive network of hydrogen bonds with R1155, Y1156, Y1184, and Y1189 as well as with conserved water molecules (Figure 1a). Less conventional interactions such as an amide-π stacking^20^ between the C-terminal glycine and W1182, and two amide NH-π bonds with W1143 and W1182, also contribute to binding. Comparison of the apo and bound states reveals a significant degree of plasticity at the binding pocket. Although the conformation of most residues is conserved, important conformational changes upon peptide binding are observed at R1155 and Y1156 (Figure 1b). Both side chains relocate to form hydrogen bonds with the peptide and bury the C-terminal di-glycine motif. Indeed, it was proposed that R1155 and Y1156 act as gatekeeper residues that move from an “open” (Figure 1c) to a “closed” (Figure 1d) conformation upon ubiquitin binding.^19^

**Figure 1.**
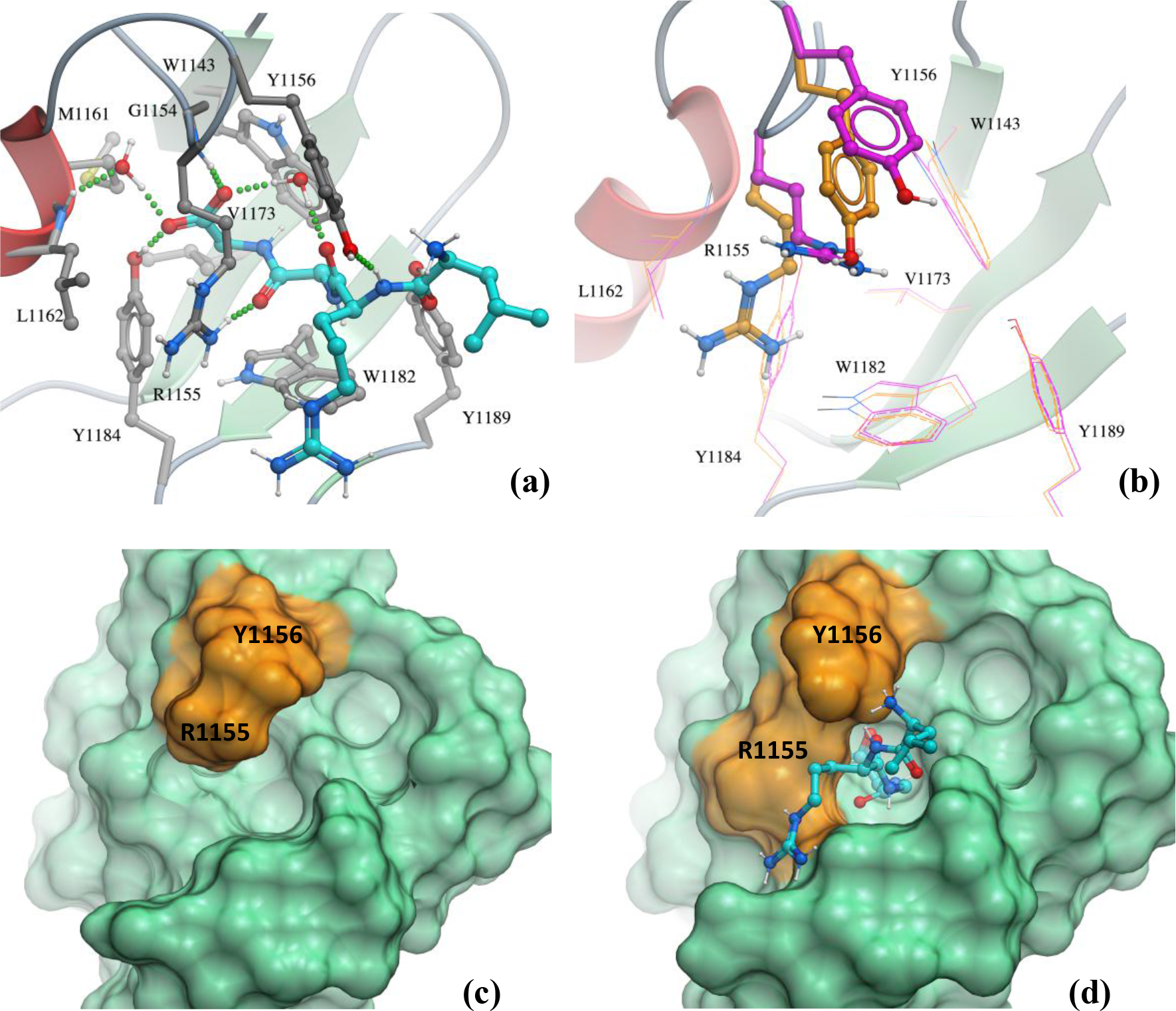
a) Crystal structure (PDB: 3GV4) of the HDAC6 ZnF-UBD domain (grey ball and sticks) in complex with a ubiquitin C-terminal pentapeptide, RLRGG (cyan ball and sticks); b) Comparison of the ZnF-UBD of HDAC6 in the apo (PDB: 3C5K, magenta) and in the peptide bound (orange) forms; c) Open (apo) conformation of HDAC6 ZnF-UBD; d) Closed (bound) conformation of HDAC6 ZnF-UBD. R1155 and Y1156 are colored in orange.

### Structure-activity relationships

To probe the chemical tractability of the ZnF-UBD, a virtual library of 1.3 million commercial compounds was docked to the ubiquitin binding pocket of HDAC6 with Glide^21^ (Schrödinger, NY), and 33 compounds (27 of which contained a carboxylate moiety) were ordered and evaluated using a fluorescence polarization (FP) peptide displacement assay. Six compounds could displace the ubiquitin peptide at micromolar concentration, and 52 readily available chemical analogs were ordered and tested, of which six were active. Overall, the hits could be grouped into clusters 1 and 2, where a carboxylic acid chain was attached to a five-or six-membered ring respectively (Table 1 and Table S1).

**Table 1.**
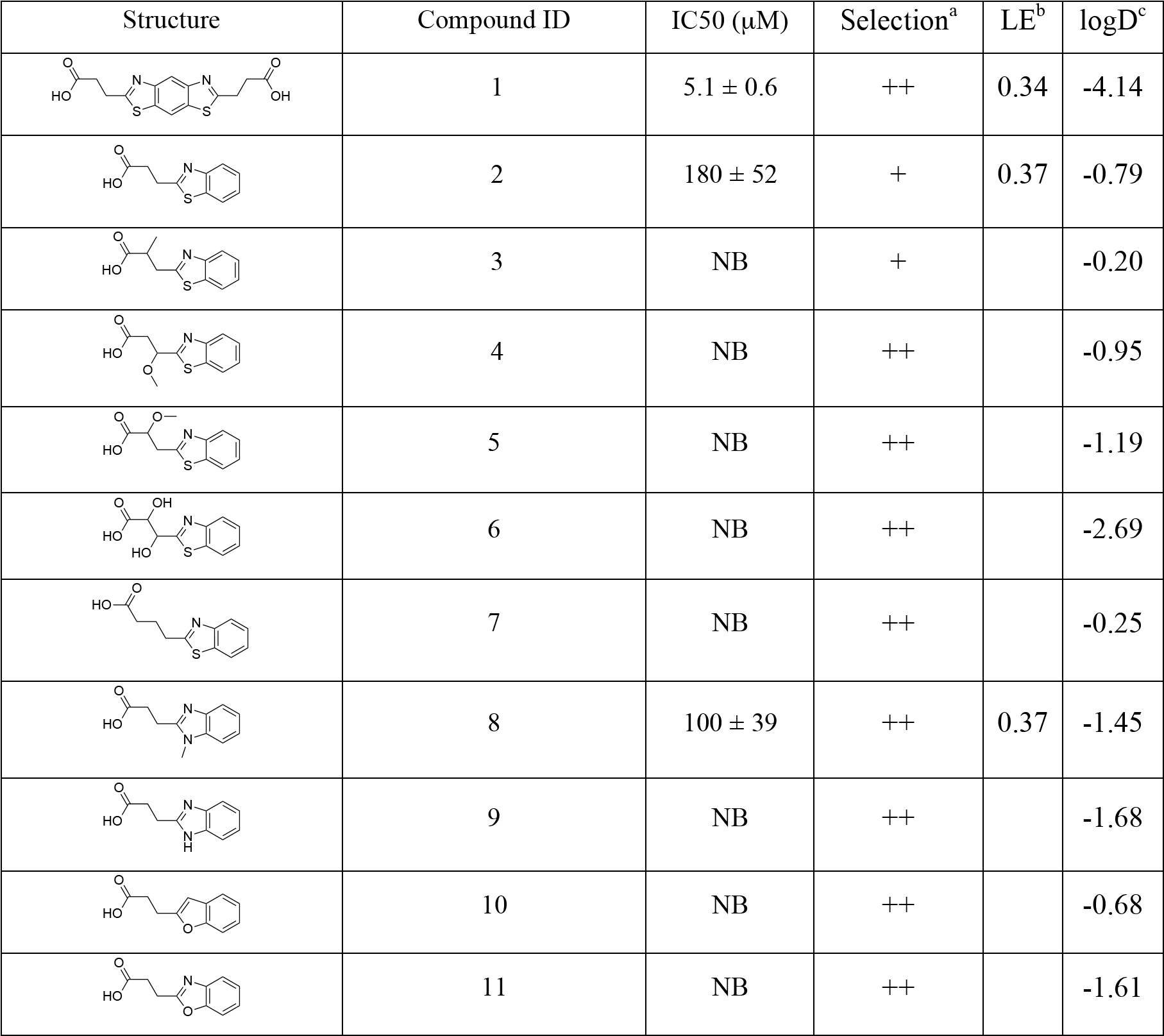
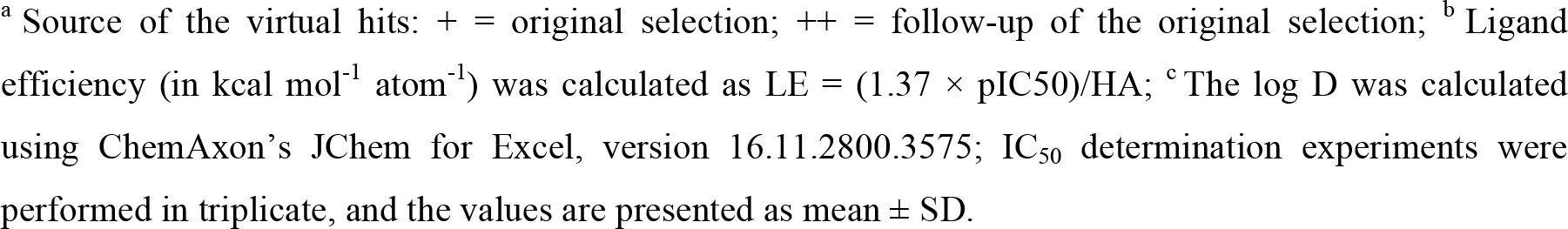
FP IC_50_ and ligand efficiency (LE) for exemplar compounds in cluster 1.

| Structure | Compound ID | IC<sub>50</sub> (μM) | Selection<sup>a</sup> | LE<sup>b</sup> | logD<sup>c</sup> |
| --- | --- | --- | --- | --- | --- |
|  | 1 | 5.1 ± 0.6 | ++ | 0.34 | -4.14 |
|  | 2 | 180 ± 52 | + | 0.37 | -0.79 |
|  | 3 | NB | + |  | -0.20 |
|  | 4 | NB | ++ |  | -0.95 |
|  | 5 | NB | ++ |  | -1.19 |
|  | 6 | NB | ++ |  | -2.69 |
|  | 7 | NB | ++ |  | -0.25 |
|  | 8 | 100 ± 39 | ++ | 0.37 | -1.45 |
|  | 9 | NB | ++ |  | -1.68 |
|  | 10 | NB | ++ |  | -0.68 |
|  | 11 | NB | ++ |  | -1.61 |

<sup>a</sup> Source of the virtual hits: + = original selection; ++ = follow-up of the original selection; <sup>b</sup> Ligand efficiency (in kcal mol<sup>-1</sup> atom<sup>-1</sup>) was calculated as LE = (1.37 × pIC<sub>50</sub>)/HA; <sup>c</sup> The log D was calculated using ChemAxon's JChem for Excel, version 16.11.2800.3575; IC<sub>50</sub> determination experiments were performed in triplicate, and the values are presented as mean ± SD.

A preliminary SAR emerged from this ensemble of molecules. Compound **1** was the most potent ZnF-UBD inhibitor, with an IC_50_ of 5.1 μM. Common to compounds **1-6** is the presence of the 3-(1,3-benzothiazol-2-yl) propanoic acid scaffold. Compound **2**, which is a substructure of compound **1**, is weaker (IC_50_ = 180 μM) but has a better ligand efficiency (LE = 0.37). Analogs of compound **2** featuring substituents on the ethylene linker (**3-6**) are inactive. Replacing the propanoic acid in **2** with a butanoic acid in **7** was not tolerated. The sulfur of compound **2** could be replaced with a hydrophobic N-methyl group in **8**. However, its analog (**9**) without the methyl group as well as other isosteres of the benzothiazole ring (**10** and **11**) are inactive.

The binding of **1** was further confirmed by surface plasmon resonance with a *K_D_* of 8 μM. We also obtained a crystal structure of **1** in complex with the ZnF-UBD of HDAC6. Although **1** may be modelled trivially in the Fo-Fc map, additional density is observed on the thiazole ring distal to the ubiquitin binding pocket, which is attributed to possible radiation damage of the ligand as described in the PDB annotation of this structure.

As shown in Figure 2a, the carboxylate of **1** overlaps well with the one from the ubiquitin peptide and forms direct or water-mediated hydrogen bonds with R1155, Y1184, L1162, and a salt bridge with R1155. Additionally, the aromatic scaffold of the compound makes π-stacking interactions with W1182 and R1155. Upon binding of **1**, R1155 and Y1156 adopt a conformation that is closer to the apo state than to the ubiquitin-bound state. The main difference is that the guanidine group of R1155 is rotated 90° to form a cation-π interaction with the aromatic ring and a hydrogen bond with the buried carboxylic acid of the ligand (Figure S1). It should be noted that the carboxylate moiety of this ligand distal to the ubiquitin binding pocket, although not fully resolved in the electron density, may make interactions with the adjacent molecule of HDAC6 in the crystal lattice which could affect its binding mode as resolved in the crystal structure. However, the close analog **2**, which is lacking this carboxylate, binds in the same orientation as **1** although is 36-fold weaker, indicating that R1155 can productively interact with the exposed carboxylate (Figure 2a).

**Figure 2.**
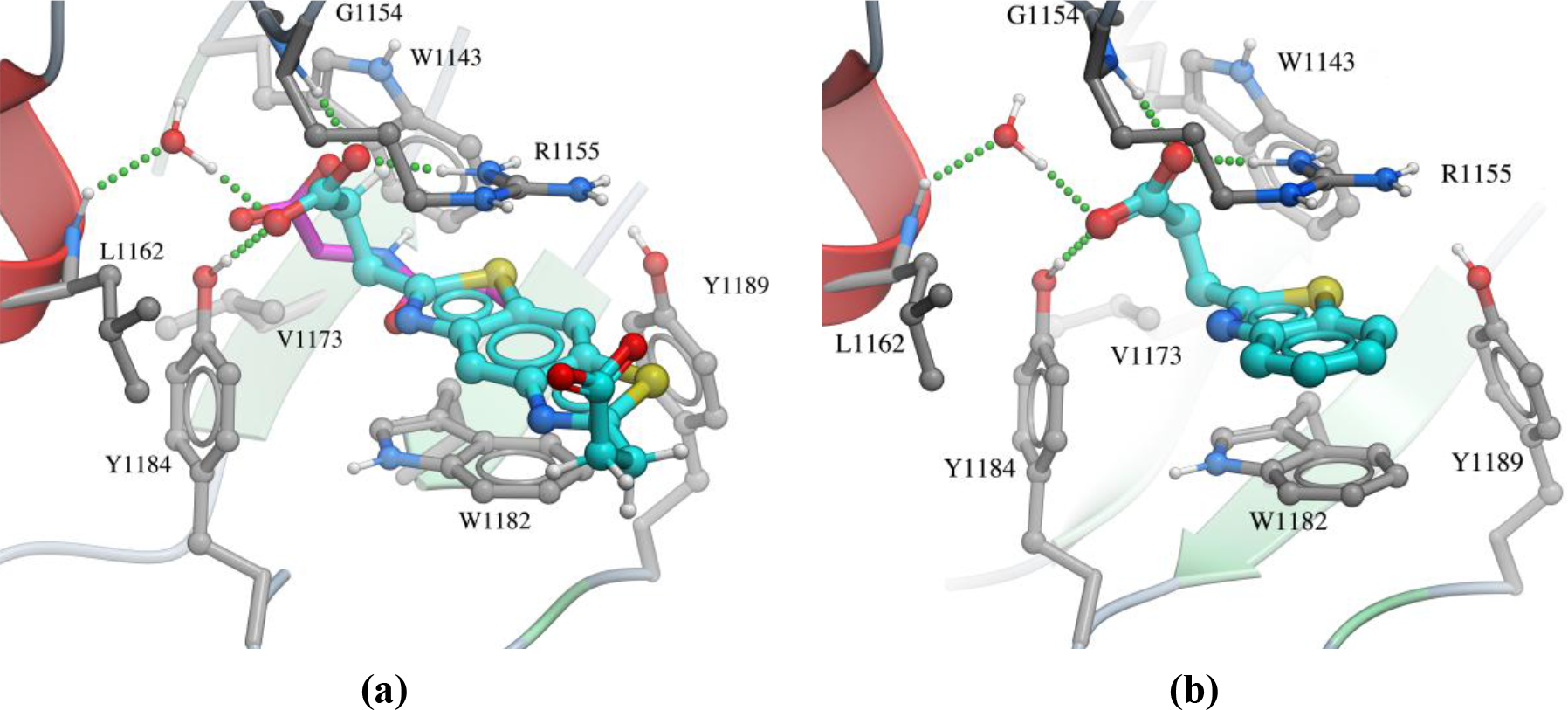
Crystal structure of HDAC6 ZnF-UBD in complex with: (a) **1** (PDB: 6CE6) and (b) **2** (PDB: 6CEF). The binding pocket residues (grey) and the inhibitors (cyan) are displayed as ball sticks and the hydrogen bonds as green dashed lines. The C-terminal glycine of the ubiquitin peptide (PDB: 3GV4) is shown as magenta sticks.

### Hit Expansion

Although compound **1** was the most potent inhibitor, it was not a favorable starting point for developing a chemical probe, as the two acidic moieties make the compound highly hydrophilic with a calculated logD and polar surface area (PSA) of -4.0 and 106 Å^2^, respectively. We focused instead on fragment **2**, which is a substructure of **1** and although weaker (IC_50_ = 180 ± 52 μM), has a slightly better ligand efficiency. We solved the crystal structure of **2** in complex with HDAC6 ZnF-UBD (Figure 2b). As expected, fragment **2** adopted a similar binding pose and recapitulated favorable interactions observed with **1**.

To circumvent the lack of close commercial analogs of **2**, several compound design strategies were employed to identify structurally related molecules of potential interest. First, we performed a substructure search focused on **2** (Figure 3a), which resulted in 111 compounds. Supported by our SAR, we envisaged that a ring expansion from the five-member ring to a six-member ring in **2** could also lead to attractive molecules. This substructure-search yielded 423 compounds (Figure 3a). Both substructure-searches were designed to retrieve compounds with one or two carbon atoms connecting the carboxylate group and the heterocycle, as our SAR showed that compounds differing only in the linker size had similar IC_50_ values (for example: **12** vs. **18**, Table S1). We also used the scaffold-hopping tool BROOD^22^ (OpenEye, CA) to identify additional chemical starting points (Figure 3b). Not surprisingly, a sulfonyl group emerged as a possible isostere of the carboxylate (Figure S2), which was an attractive mechanism to improve the physicochemical properties of the compound, at the risk of losing electrostatic interactions with R1155. A substructure search for analogs of **2** with the sulfonyl group yielded 315 commercial compounds (Figure 3b).

**Figure 3.**
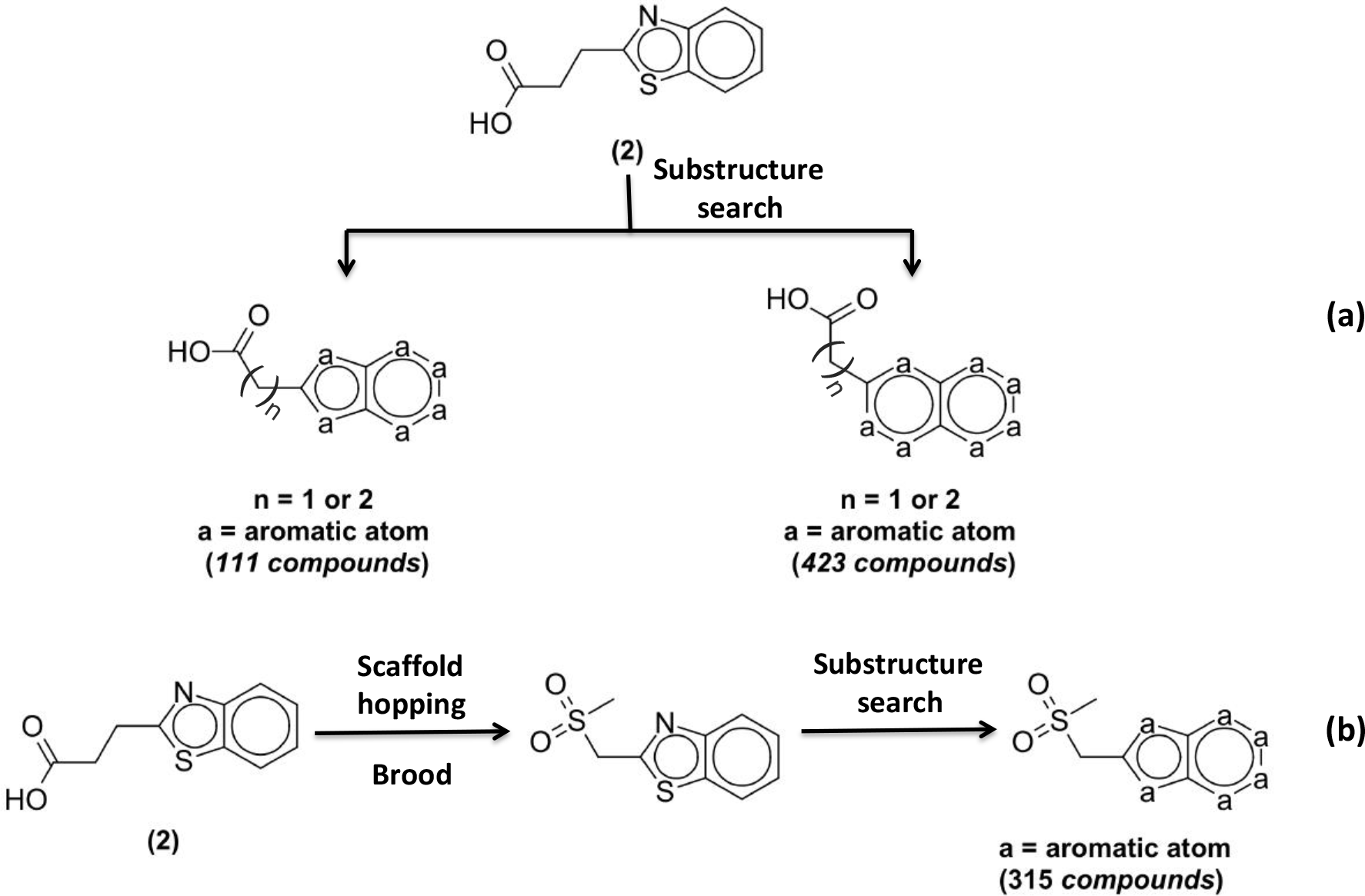
Design strategy for the hit expansion: (a) Substructure search and ring expansion of analogs of **2**; (b) Scaffold-hopping followed by substructure search.

The resulting library of 849 compounds was then docked to the structure of HDAC6 ZnF-UBD in complex with **1** using Glide^21^ (Schrodinger, NY), with three hydrogen bond constraints at the side chain OH of Y1184, the NH backbone and the side chain of R1155. After careful visual inspection, 100 compounds that satisfied the hydrogen bond constraints and also formed a π-stacking interaction with W1182 and R1155 were selected and rescored using the molecular mechanics generalized Born surface area (MM/GBSA) method with AMBER 12.^23^ Upon visual inspection, 13 compounds were ordered, and tested using a fluorescence polarization peptide displacement assay. Of the 13 compounds ordered, 10 displaced the ubiquitin peptide at micromolar concentration (Table 2).

**Table 2.**
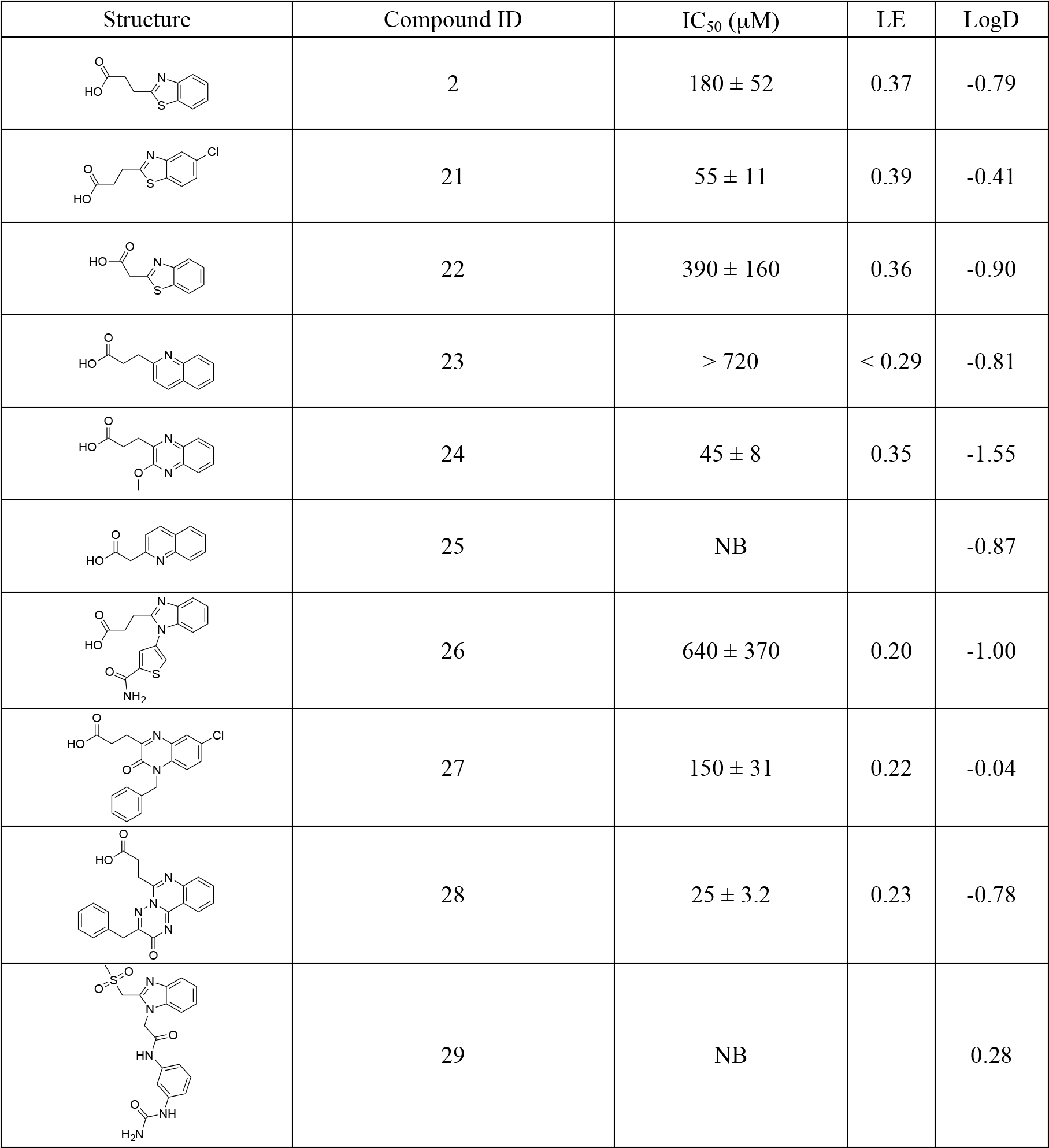
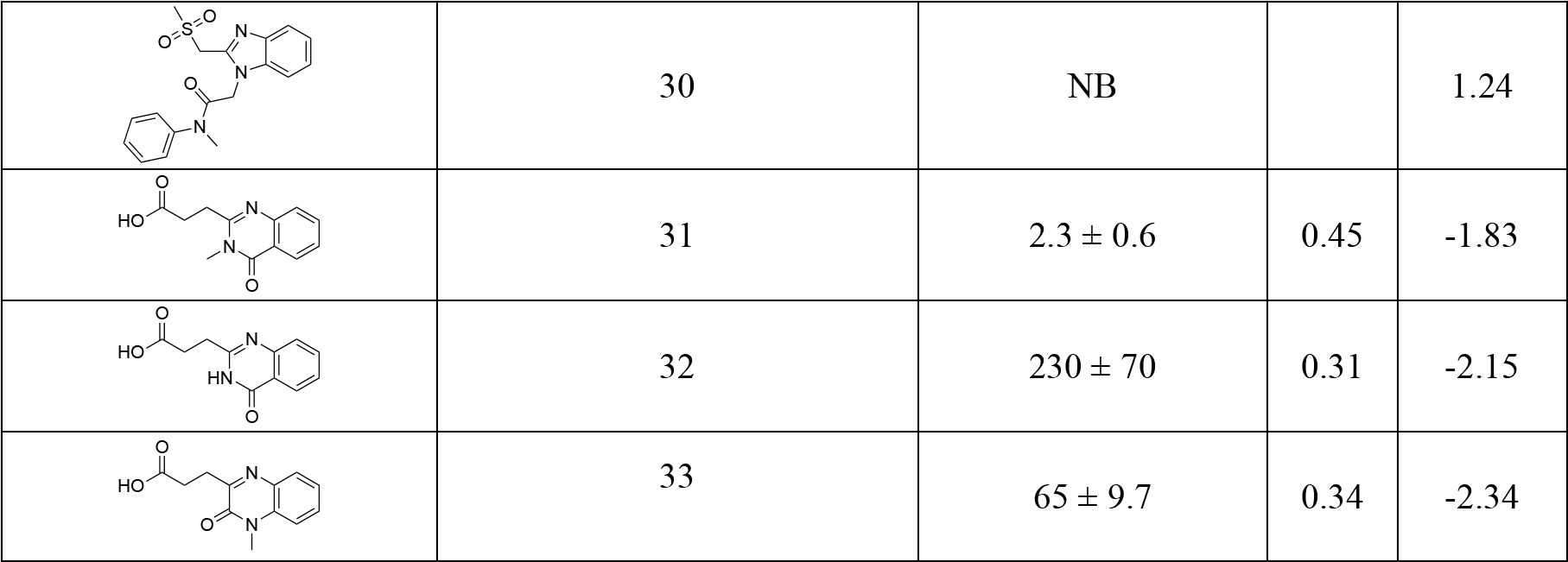
Follow-ups of **2** are shown, and annotated with IC_50_ values that were determined using a fluorescence polarization assay.

| Structure | Compound ID | IC<sub>50</sub> (μM) | LE | LogD |
| --- | --- | --- | --- | --- |
|  | 2 | 180 ± 52 | 0.37 | -0.79 |
|  | 21 | 55 ± 11 | 0.39 | -0.41 |
|  | 22 | 390 ± 160 | 0.36 | -0.90 |
|  | 23 | > 720 | < 0.29 | -0.81 |
|  | 24 | 45 ± 8 | 0.35 | -1.55 |
|  | 25 | NB |  | -0.87 |
|  | 26 | 640 ± 370 | 0.20 | -1.00 |
|  | 27 | 150 ± 31 | 0.22 | -0.04 |
|  | 28 | 25 ± 3.2 | 0.23 | -0.78 |
|  | 29 | NB |  | 0.28 |
|  | 30 | NB |  | 1.24 |
| | 31 | $2.3 \pm 0.6$ | 0.45 | -1.83 |
| | 32 | $230 \pm 70$ | 0.31 | -2.15 |
| | 33 | $65 \pm 9.7$ | 0.34 | -2.34 |

The data provided further insight into the structure activity relationships of this chemical series. Adding a chlorine at position C5 of the benzothiazole ring resulted in a 3-fold gain in potency (compound **21)**, but removing a methylene from the acidic side chain was detrimental (compound **22**). Compound **24**, with a 3-methoxyquinoxaline scaffold inhibited HDAC6 ZnF-UBD with an IC_50_ of 45 μM. Comparing **2** vs. **22**, and **23** vs. **25** indicated that a propionic was preferred over an acetic acid. Compounds **26-28** were selected to extend interactions beyond the ubiquitin C-terminus binding pocket, of which **28** inhibited the protein with an IC_50_ of 25 μM (LE = 0.23). Finally, the two sulfonyl compounds (**29** and **30**) were inactive.

The quinazolinone scaffold of **31** represented a significant breakthrough in the potency and ligand efficiency of this chemical series with an IC_50_ of 2.3 μM by FP (LE = 0.45), and a *K_D_* of 1.5 μM determined by SPR. Removing the critical N-methyl group at the amide moiety (compound **32)** or shifting its position as in the regioisomer **33**, resulted in a 100-or 28-fold loss in potency, respectively.

To investigate the structural basis for the relative potencies, all 13 compounds were soaked into crystals of HDAC6 ZnF-UBD, and seven crystal structures were obtained with resolutions ranging from 1.55 to 1.75 Å. The docked poses were confirmed for six of the seven structures (RMSD 0.3 to 1.9 Å – Figure S3) and recapitulated the general binding mode previously observed with fragment **2**, including an anchored carboxylate warhead, salt bridges and a cation-π interaction with R1155, as well as a π- stacking interaction with W1182 (Figures 4 and 5).

**Figure 4.**
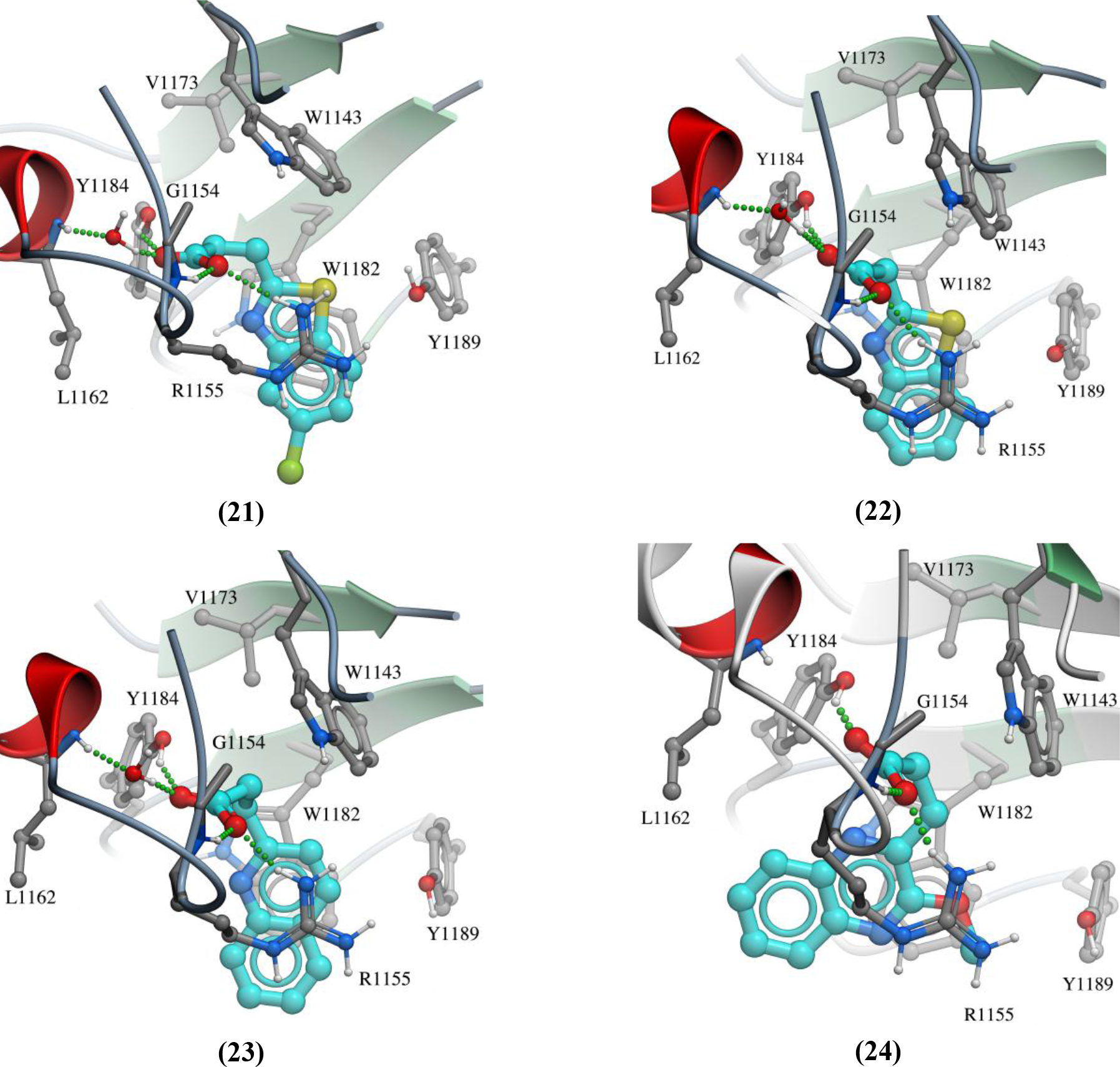
Crystal structures of four fragments in complex with HDAC6 ZnF-UBD: **21** (PDB: 5KH3); **22** (PDB: 6CE8); **23** (PDB: 6CEA); **24** (PDB: 6CEC). The compounds (cyan) and the amino acid residues (grey) are displayed as ball and sticks. Hydrogen bonds are displayed as green dashed lines.

**Figure 5.**
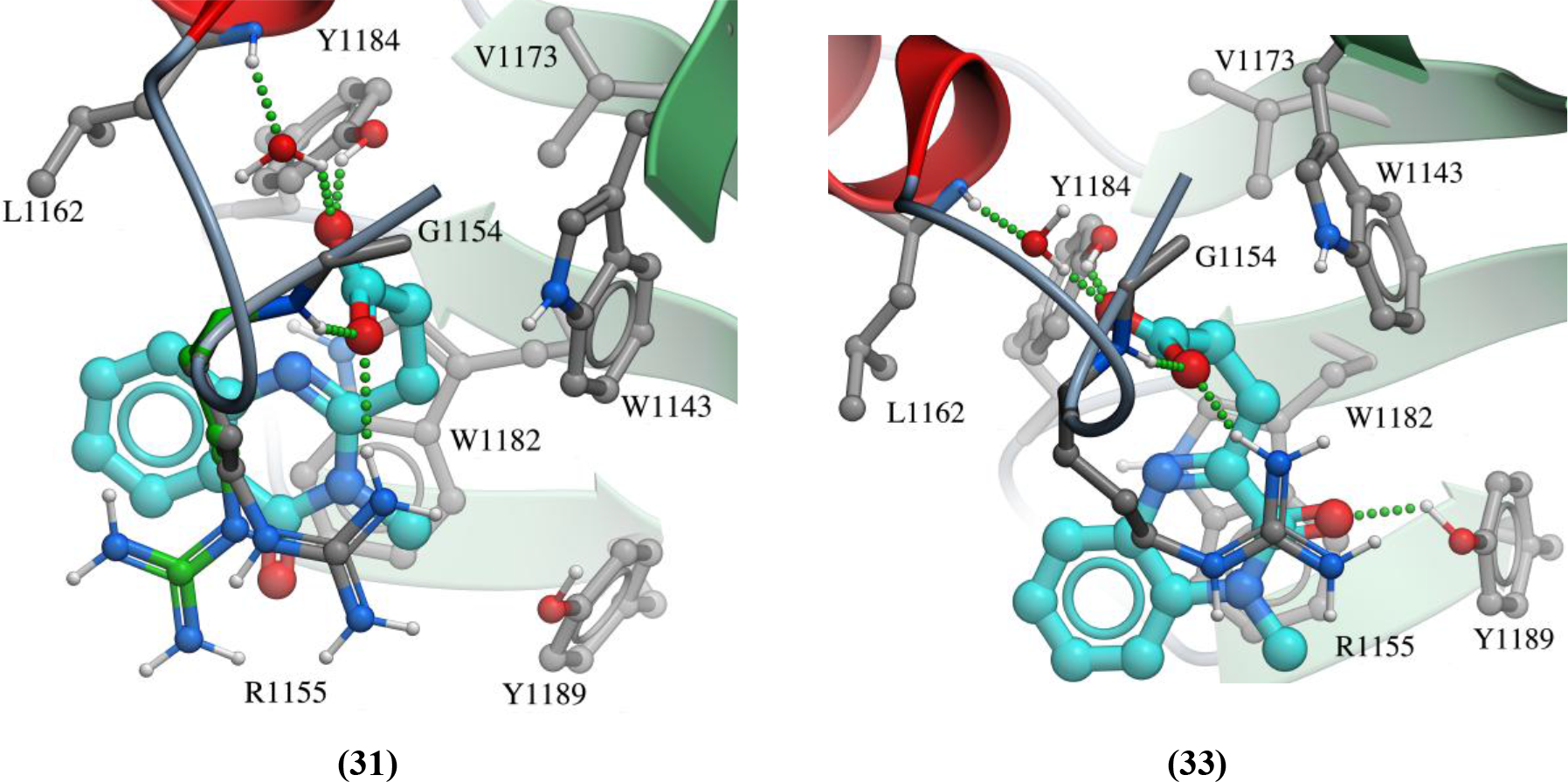
Crystal structures of **31** (PDB: 6CED) and **33** (PDB: 6CEE) in complex with HDAC6 ZnF-UBD. The compounds (cyan) and the amino acid residues (grey) are displayed as ball and sticks. Hydrogen bonds are displayed as green dashed lines. The lower occupancy of R1155 in **31** is displayed as green ball and sticks.

As expected, **21** binds in the same conformation as **2**; the chlorine group pointed towards the solvent and is probably extending or strengthening π-π interactions with neighboring side-chains (Figure 4). Fragment **22**, featuring an acetic instead of a propionic acid, preserved the interactions observed for **2** and the benzothiazole rings of the two compounds overlapped perfectly (Figure 4). Based on the crystal structures, the increased potency of **2** vs. **22** probably reflects a reduced conformational strain associated with a longer carboxylic chain.

While the amino acid side-chains lining the inhibitor binding pocket adopted identical conformations in the previous four structures (Figure 4), R1155 was also found in an alternate, lower-occupancy conformation in complex with **31**, where the guanidine group was flipped ~180° (Figure 5). The critical methyl group of **31** occupies a cavity formed by the polar guanidine of R1155 and the hydroxyl group of Y1189, as well as the partially hydrophobic side-chain of W1182. The combined loss of hydrophobic interactions with W1182 and electronic rearrangement at the aromatic system are probably responsible for the 100-fold decrease in potency upon removal of this methyl group (compound **32**). The gain of a hydrogen bond with the carbonyl group of the regioisomer **33** (Figure 5) partially mitigates this loss (28-fold drop in potency).

### A bicyclic ring system and a carboxylate are essential for binding

The complex of HDAC6 ZnF-UBD with **24** was also solved: the methoxy group of the fragment points towards the side chain of Y1189, but does not form productive interactions with this residue (Figure 4). Only one of the two aromatic rings of **24**, makes π-stacking and cation-π interactions with W1182 and R1155, respectively, while both rings were well positioned to make these interactions in the weaker compound **23** (Figure 4). This raised the possibility that these fragments could be simplified to a monocyclic ring system. To test this hypothesis we synthesized compound **34**, the single-ring analog of **24** (Table 3). No detectable binding was observed, supporting the notion that a fused bicyclic aromatic ring system is necessary.

**Table 3.**
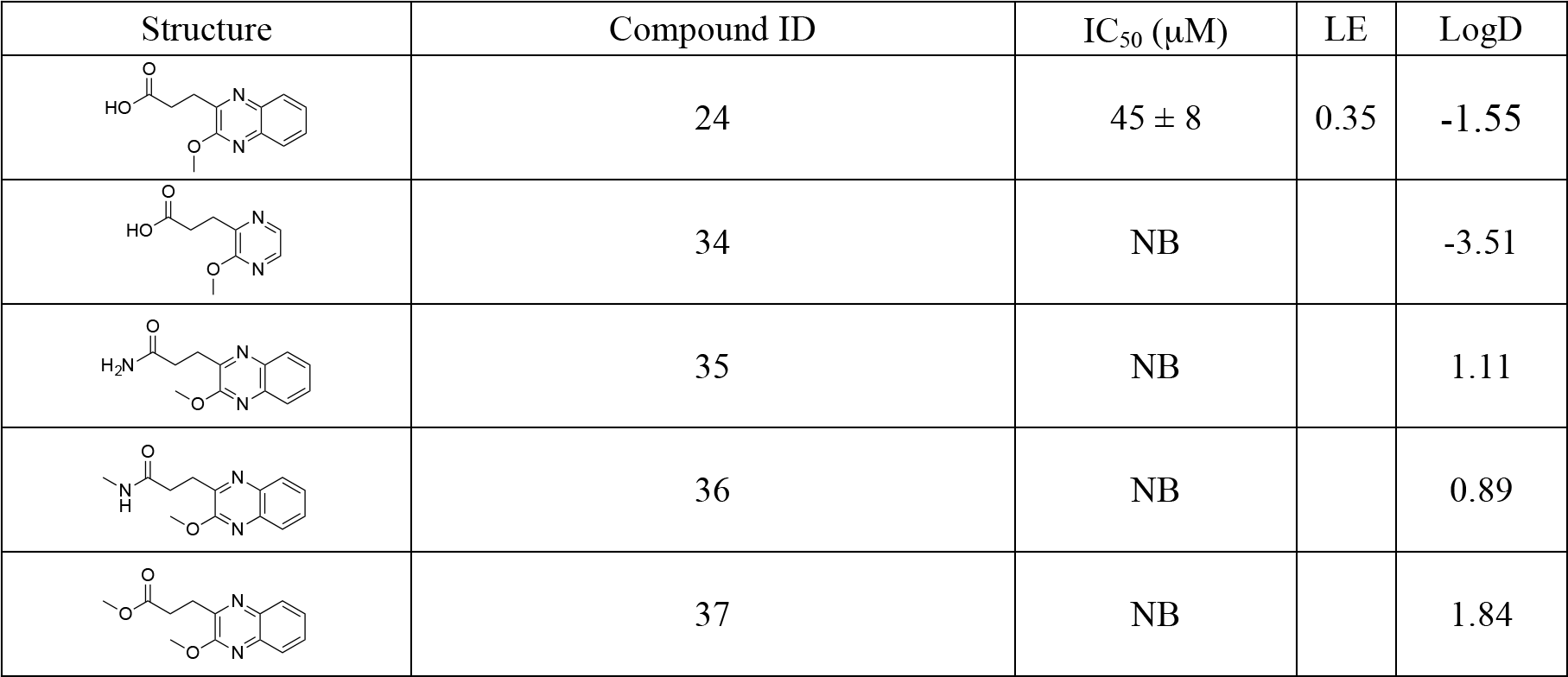
Structure, IC_50_ and LE values of compounds used to investigate the importance of the ring size and of the carboxylate group.

| Structure | Compound ID | IC<sub>50</sub> (μM) | LE | LogD |
| --- | --- | --- | --- | --- |
|  | 24 | 45 ± 8 | 0.35 | -1.55 |
|  | 34 | NB |  | -3.51 |
|  | 35 | NB |  | 1.11 |
|  | 36 | NB |  | 0.89 |
|  | 37 | NB |  | 1.84 |

Replacement of the carboxylate with a primary or secondary amide (**35** and **36**, respectively) or an ester group (**37**) was not tolerated, indicating that the conserved network of direct and water-mediated interactions between the carboxylate and R1155, Y184, and L1162 is essential (Figure 4, Figure 5, Table 3).

### Conformational rearrangements can open the door to a side pocket

Crystal structures of HDAC6 ZnF-UBD feature a well-defined cavity juxtaposed to the binding pocket exploited by the compounds presented here. This secondary pocket is separated from the primary pocket by a wall composed of Y1189 and R1155 (Figure 6a). Interestingly, in the structure in complex with **28**, this wall disappears, raising the possibility of simultaneously exploiting both cavities (Figure 6b). Here, R1155 adopts an unique conformation (Figure S4) where it flips away from the secondary pocket to stack with a third fused ring of the central core (absent from other inhibitors) of **28**, opening the door between the two binding pockets (Figure 6c).

**Figure 6.**
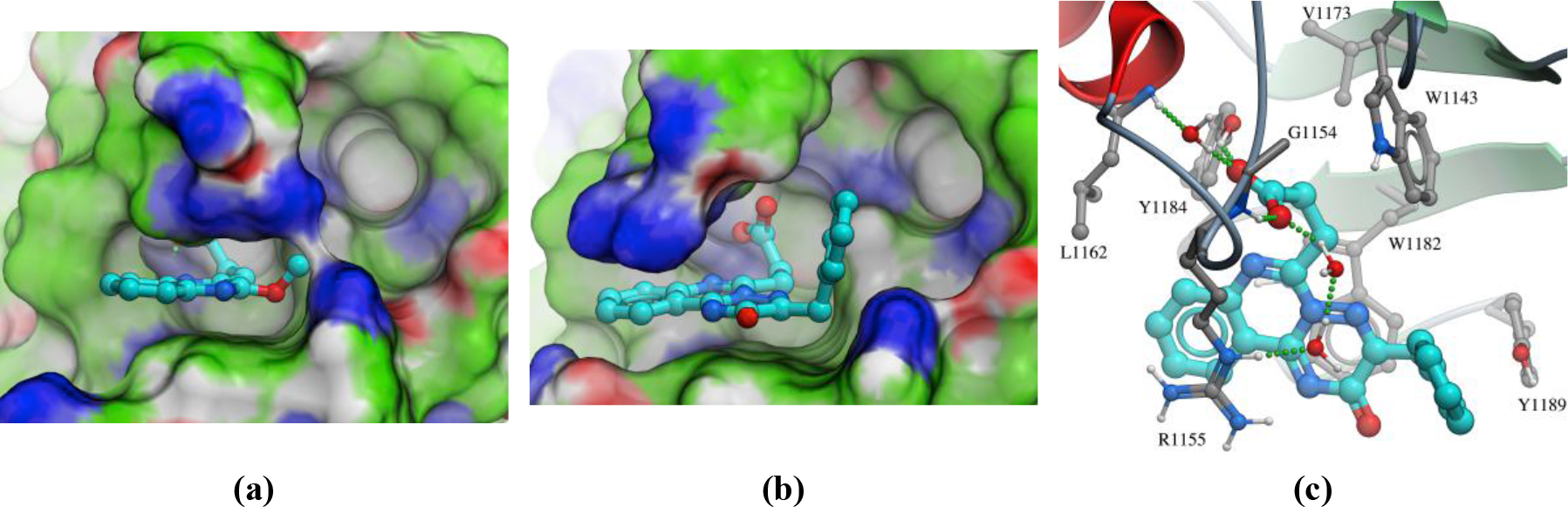
Crystal structure of HDAC6 ZnF-UBD in complex with **24** (PDB: 6CEC) (a) and **28** (PDB: 5WBN) (b, c), respectively. The compound (cyan) and the amino acid residues (grey) are displayed as ball and sticks. Hydrogen bonds are displayed as green dashed lines. The surface is colored by binding property: white = neutral; green=hydrophobic; blue=hydrogen bonding donor potential; red=hydrogen bond acceptor potential.

We have recently reported weak fragments occupying the secondary pocket and a weak inhibitor occupying both pockets suggesting that it is ligand-able.^18^ However, the π-stacking of the heterocyclic core with the R1155 and W1182 is disrupted in this inhibitor resulting in weak affinity. Compound **28** is not a favorable scaffold due to chemical liabilities, but the crystal structure and associated IC_50_ of 25 μM demonstrates the feasibility of grow the inhibitor to occupy both pockets without disrupting the key interactions in the main pocket as a promising strategy to improve potency.

### Compound 31 displaces the RLRGG peptide from full length HDAC6

We also used a fluorescence polarization assay to test whether fragment **31** would bind to the ZnF-UBD domain in the context of full length HDAC6, a pre-requisite for the future development of a cell-active chemical probe. Our result shows that **31** also displaces the C-terminal ubiquitin peptide from full length HDAC6 with an IC_50_ of 3.5 ± 0.9 μM (Figure 7). This value showed no significant difference from 2.3 ± 0.6 μM obtained with the ZnF-UBD domain alone, and shows that **31** does bind to HDAC6 in the context of the full-length protein.

**Figure 7.**
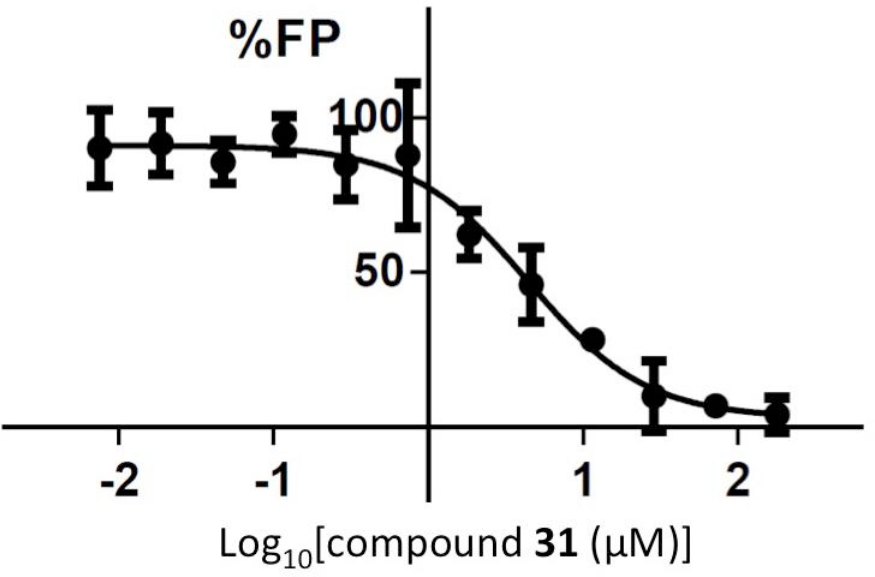
FP competition assay using increasing concentrations of compound **31**, a FITC-labelled RLRGG peptide (50 nM) and full length HDAC6 (1 μM). An IC_50_ of 3.5 ± 0.9 μM was obtained from the average of three independent measurements.

## Conclusion

Here, we report the successful attempt to target the ZnF-UBD of HDAC6 with small molecules selected by virtual screening. SAR analysis indicates that a carboxylate warhead, linked to a fused bicyclic aromatic ring system that can stack with neighboring tryptophan and arginine side-chains are important chemical features for activity. Inhibitor-induced conformational remodeling at the binding site can open-up a channel towards a ligand-able side-pocket that may be exploited to increase potency. We believe that the chemical inhibitors presented here and accompanying crystal structures will inform the discovery of a novel class of HDAC6-selective chemical probes.

## Experimental Section

### Primary virtual screening

Ligprep^24^ (Schrodinger, NY) was used to convert to 3D a diverse virtual library of 1,280,604 drug-like compounds from commercial vendors (protonation states at pH 7.3 +/- 0.2 were generated with Epik). The resulting library was docked to the ubiquitin-binding pocket of HDAC6 (PDB code 3GV4) with Glide HTVS and the top 10% scoring compounds were re-docked with Glide SP (Schrodinger, NY).^21^ The process was conducted twice independently, with or without a water molecule at the bottom of the binding pocket (forming a hydrogen-bond with the main-chain of C1153 and L1162). The 213 compounds scoring better than -7.0 in the presence of the water molecule and the 100 compounds scoring better than -6.5 in the absence of water were selected for visual inspection, from which 40 compounds were subjectively picked, based on chemical scaffold and binding pose, and 33 were in stock and ordered.

### Substructure search

All substructure searches were performed using the software FILTER^25^ (OpenEye, CA) against ~22 million commercial compounds downloaded from the ZINC database.^26^ LigPrep^24^ was used to prepare the ligands using default settings.

### Scaffold hopping

BROOD^22^ (OpenEye, CA) was used to select potential isosteric groups of the carboxylic acid using the fragment [R1]CCC(=O)[O-] as the query. BROOD generates analogs of a compound by replacing selected fragments (scaffolds) in the molecule with fragments that have similar shape and electrostatics. The sulfonyl group was the best group to match the query fragment in terms of shape, electrostatics, and attachment geometry.

### Ligand docking

The library designed by the substructure search and scaffold hopping exercise was docked to the HDAC6 Zn-UBD structure in complex with **1** using Glide with default settings. Three hydrogen bond constraints at the side chain OH of Y1184, the NH backbone and the side chain of R1155 were used in the docking calculations.

### Molecular mechanics (MM/GBSA)

The docked poses of the ligands in complex with HDAC6 Zn-UBD were rescored using the MM/GBSA method.^27^ Missing atoms and hydrogens were added using the tleap program in Amber12 software.^23^ Amber ff99SB^28^ force field and GAFF force field^29,30^ were assigned respectively to amino acid residues and to the ligands. The GAFF parameters were generated using antechamber and parmchk program, and partial atomic charges were derived using AM1-BCC methodology.^31^ Each complex structure was solvated in a TIP3P water cubic box extending at least 10 Å from the complex, and then neutralized by the addition of appropriate number of monovalent counter-ions.

### Cloning, Protein Expression and Purification

DNAs encoding HDAC6^1109-1213^ and HDAC6^1109-1215^ were subcloned into a modified pET28 vector encoding a thrombin cleavable (GenBank EF442785) N-terminal His6-tag (pET28-LIC) and HDAC6^1109-1215^ was also sub-cloned into a modified pET28 vector encoding an N-terminal AviTag for in vivo biotinylation and a C-terminal His6-tag (p28BIOH-LIC) using a ligation-independent InFusion™ cloning kit (ClonTech) and verified by DNA sequencing. Proteins were over-expressed in BL21 (DE3) Codon Plus RIL E. coli (Agilent). Cultures were grown in M9 minimal media supplemented with 50 μM ZnSO4 and also 10 μg/mL biotin for HDAC6^1109-1215^ p28BIOH-LIC expression. Expression cultures were induced using 0.5 mM IPTG overnight at 15 °C. Proteins were purified using nickel-nitrilotriacetic acid (Ni-NTA) agarose resin (Qiagen) and the tag was removed by thrombin for HDAC6^1109-1213^ pET28-LIC. Uncleaved proteins and thrombin were removed by another pass with Ni-NTA resin. Proteins were further purified using gel filtration (Superdex 75, GE Healthcare). The final concentrations of purified proteins were 5-10 mg/mL as measured by UV absorbance at 280 nm.

DNA encoding HDAC6^1-1215^ was cloned into a modified derivative pFastBac Dual vector (Invitrogen) encoding an N-terminal AviTag and a C-terminal His6 tag (pFBD-BirA). Cloning was completed using a ligation-independent InFusionTM cloning kit (ClonTech) and verified by DNA sequencing. HDAC6^1-1215^ was over-expressed in sf9 insect cells. Cultures were grown in HyQ SFX Insect Serum Free Medium (Fisher Scientific) to a density of 4×10^6^ cells/mL and infected with 10 mL of P3 viral stock media per 1 L of cell culture. Cell culture medium was collected after 4 days of incubation in a shaker. Protein was purified using nickel-nitrilotriacetic acid (Ni-NTA) agarose resin (Qiagen). Following overnight dialysis, protein was further purified using anion exchange (HiTrap Q HP, GE Healthcare) and gel filtration (Superdex 75, GE Healthcare). The final concentration of purified HDAC6^1-1215^ was 3.5 mg/mL as measured by UV absorbance at 280 nm.

### Crystallization

The apo crystal form of HDAC6^1109-1215^ (PDB ID: 3C5K) was previously reported.^19^ These crystals can be used to seed for the crystal form amenable to soaking. Diluting 1 μL crystal suspension 1:10,000 with mother liquor and vortexing the sample vigorously yielded a seed mix. HDAC6^1109-1213^ can be crystallized in 2 M Na formate, 0.1 M Na acetate pH 4.6, 5 % ethylene glycol in 5:4:1 3.5 mg/mL protein, mother liquor and seed mix per drop. HDAC6^1109-1213^ crystals were soaked by adding 5 % (v/v) of a 200 mM or 400 mM DMSO-solubilized stock of compounds to the drop for 2 hours prior to mounting and cryo-cooling.

### Data Collection, Structure Determination and Refinement

X-ray diffraction data for HDAC6^1109-1213^ cocrystals were collected at 100K at Rigaku FR-E Superbright home source at a wavelength of 1.54178 Å. All datasets were processed with XDS^32^ and Aimless^33^. Models were refined with cycles of COOT^34^ for model building and visualization, with REFMAC^35^, for restrained refinement and validated with MOLPROBITY^36^.

### Surface Plasmon Resonance (SPR)

Studies were performed using a Biacore T200 (GE Health Sciences). Approximately 5000 response units (RU) of biotinylated HDAC6^1109-1215^ were coupled onto one flow cell of a SA chip as per manufacturer’s protocol, and an empty flow cell used for reference subtraction. 2-fold serial dilutions of compounds were prepared in 10 mM HEPES pH 7.4, 150 mM NaCl, 0.005 % (v/v) Tween-20, 1 % (v/v) DMSO. KD determination experiments were performed using single-cycle kinetics with 30 s contact time, 30 μL/min flow rate at 20 °C. KD values were calculated using steady state affinity fitting and the Biacore T200 Evaluation software.

### Fluorescence Polarization (FP) Displacement Assay

#### RLRGG Peptide andHDAC6 ZnF-UBD

All experiments were performed in 384-well black polypropylene PCR plates (Axygen) in 10 μL volume. Fluorescence polarization (FP) was measured using a BioTek Synergy 4 (BioTek) at excitation and emission wavelengths were 485 nm and 528 nm, respectively. In each well, 9 μL compound solutions in buffer containing 10 mM HEPES pH 7.4, 150 mM NaCl, 1 % (v/v) DMSO were serially diluted. 1 μL 30 μM HDAC6^1109-1215^ and 500 nM N-terminally FITC-labelled RLRGG were then added to each well. Following 1 min centrifugation at 250 g, the assay was incubated for 10 min before FP analysis. Previous assay optimization with titration of protein concentration at 50 nM peptide showed these conditions gave ~85 % maximum FP.

#### RLRGG Peptide and Full Length HDAC6 Assay

Experiments were performed in a total volume of 10 μL in a 384-well black polypropylene PCR plate (Axygen). FP was measured using Synergy 4 (BioTek) after 10 min of incubation. The excitation and emission wavelengths were 485 nm and 528 nm, respectively. Each well contained 9 μL of compound 31 (200 μM) in buffer containing 20 mM Hepes pH 7.4, 150 mM NaCl, 0.005% (v/v) Tween-20, 0.1% (v/v) DMSO serially diluted. 1 μL of 10 μM HDAC6^1-1215^ and 500 nM N-terminally FITC-labelled RLRGG was then added to each well. The plate was centrifuged at 150 g for 1 min before FP analysis.

### Chemistry and Compound Purity

Compounds **21**, **24**, **31-33**, and **34-37** were synthesized according to the procedures described in the Supporting Information. All commercial and synthesized compounds tested in vitro were ≥ 95% pure. Purity was determined by analytical HPLC on a Agilent 1100 series instrument equipped with a Phenomenex KINETEX column (50.0 mm × 4.6 mm, C18, 2.6 μM) at 25 °C. A linear gradient starting from 5% acetonitrile and 95% water to 95% acetonitrile and 5% water over 4 minutes followed by elution at 95% acetonitrile and 5% water was employed. Formic acid (0.1%) was added to all solvents.

### Accession Codes

Atomic coordinates and experimental data were deposited in the Protein Databank, and will be released immediately. HDAC6 ZnF-UBD co-crystal structure PDB codes: **1** (6CE6), **2** (6CEF), **22** (6CE8), **23** (6CEA), **24** (6CEC), **28** (5WBN), **31** (6CED), **33** (6CEE).

## Acknowledgments

The SGC is a registered charity (number 1097737) that receives funds from AbbVie, Bayer Pharma AG, Boehringer Ingelheim, Canada Foundation for Innovation, Eshelman Institute for Innovation, Genome Canada through Ontario Genomics Institute [OGI-055], Innovative Medicines Initiative (EU/EFPIA) [ULTRA-DD grant no. 115766], Janssen, Merck KGaA, Darmstadt, Germany, MSD, Novartis Pharma AG, Ontario Ministry of Research, Innovation and Science (MRIS), Pfizer, São Paulo Research Foundation-FAPESP, Takeda, and Wellcome [106169/ZZ14/Z]. IF and MM are supported by NSERC CREATE grant 432008-2013. Andrei Yudin (University of Toronto) is thanked for analytical HPLC access. We are grateful to OpenEye Scientific Software, Inc. for providing us with an academic license for their software.

## Table of Contents graphic

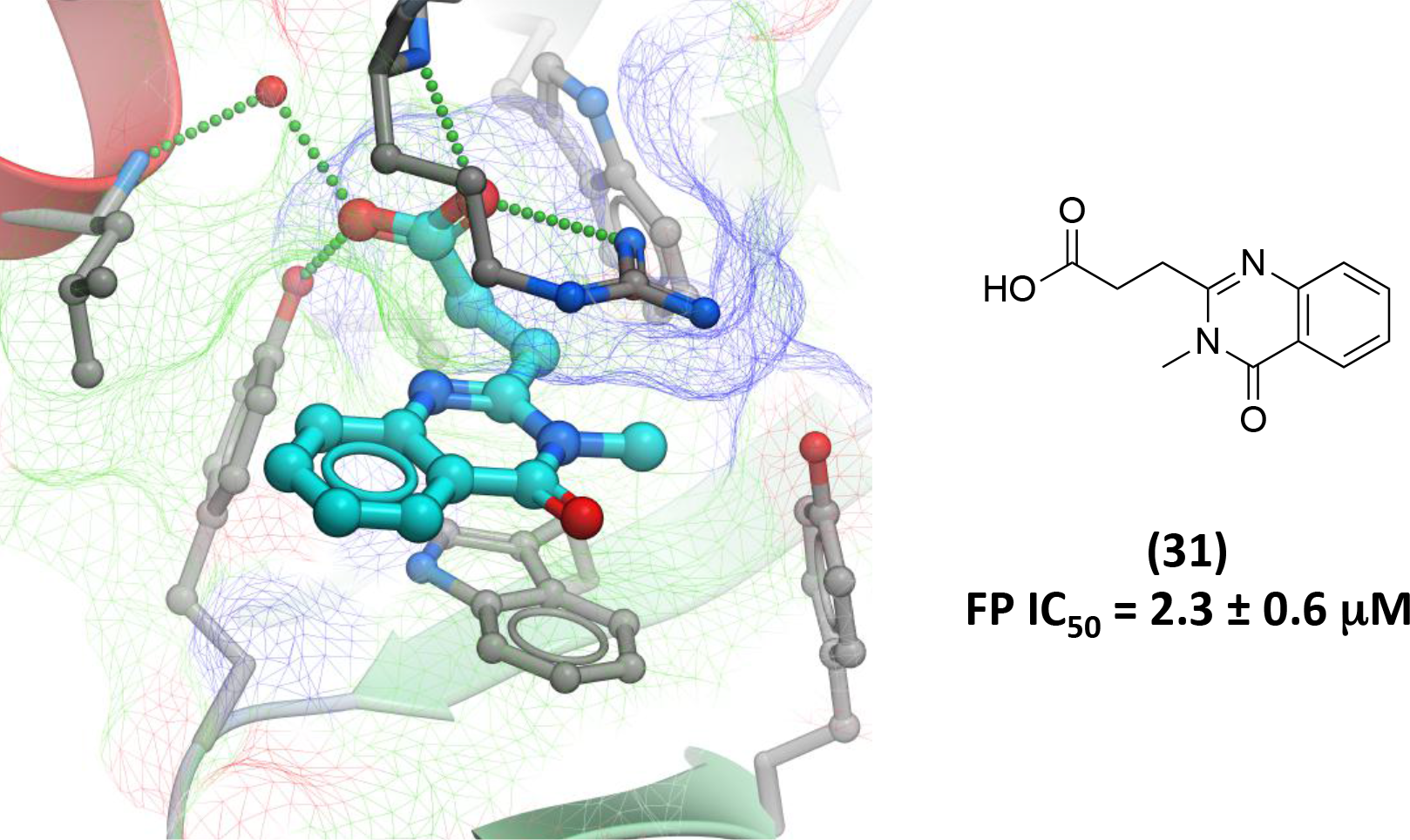

